# Additive manufacturing of PEDOT:PSS electrodes on collagen substrates for soft and bioactive electronics

**DOI:** 10.64898/2026.05.08.723335

**Authors:** Tianran Liu, Jae Park, Somtochukwu S. Okafor, Sandra K. Montgomery, Anna P. Goestenkors, Barbara A. Semar, Riley M. Alvarez, Cayleigh P. O’Hare, Yuqing Wu, Justin S. Yu, Cielo J. Vargas Espinoza, Alexandra L. Rutz

## Abstract

Traditional bioelectronic devices are limited by poor biointerfacing due to their substantial mismatch in mechanical and biochemical properties. In tissue engineering, soft and bioactive materials support biointegration by harnessing or mimicking the natural extracellular matrix (ECM). Building bioelectronic devices from ECM should improve their biointegration, yet there are limited methods to fabricate them due to current manufacturing approaches. An additive manufacturing strategy is presented here for collagen-based bioelectronic interfaces that integrates conducting polymer electrodes with ECM-based substrates or encapsulation layers. Addition of poly(ethylene glycol) diglycidyl ether (PEGDE) to poly(3,4-ethylenedioxythiophene):poly(styrene sulfonate) (PEDOT:PSS) colloidal dispersions enables direct extrusion-based patterning under mild conditions compatible with collagen substrates, and forms aqueous stable and highly conducting printed patterns (2788 S m⁻¹). The resulting interfaces maintain stable electrochemical performance over 7 days in physiological environments, and support primary human cell adhesion, viability, and proliferation across both material regions. A sacrificial patterning strategy using 3D printed cacao butter further enables spatial control of collagen encapsulation. This approach establishes a framework for fabricating functional bioelectronic devices based on ECM to further enhance device biointerfaces for tissue models and implantable systems.

## 1. Introduction

Electrical phenomena play a fundamental role in biological processes that enable living systems to form, repair, and operate (e.g., stem cell differentiation^1^, tissue regeneration^1^, neuronal signaling and muscle actuation^2^). To investigate and modulate these crucial processes, researchers and clinicians rely on bioelectronic devices to monitor electrical activity or deliver controlled stimulation^3,4^. To best serve these purposes, ideal bioelectronics should seamlessly integrate with biological systems. Yet, tissue integration of conventional bioelectronics is inherently challenged by the biointerfacial materials^3,5,6^. Materials traditionally comprising electronic devices are often dry, stiff, and biologically inert. These conventional materials include conductors like gold and silicon as well as substrates and encapsulation layers like glass, silicone, polyimide, and SU-8^3,5,6^. In contrast, the typical cellular microenvironment consists of hydrated, soft extracellular matrices (ECM) that actively participate in cellular signaling. As a result of this mismatch in material properties, both *in vitro* and *in vivo* bioelectronic interfaces suffer from suboptimal integration. *In vitro,* these mismatches in cellular microenvironments can result in abnormal cell behaviors^7^, limiting the physiological relevance and translational value of experimental findings^8^. *In vivo*, the failure to integrate with the host leads to a foreign body response and fibrotic encapsulation, impairing sensing and stimulation^9,10^.

To improve biologic-electronic integration, existing approaches have broadly focused on either mechanical or biochemical properties of the biointerface. Mechanical strategies aim to engineer devices that better resemble soft tissue. Such approaches may include reducing device stiffness via structural design (e.g., mesh^11^, thin film^12^, and fiber electrodes^13^) or incorporation of soft materials^14–18^. In parallel, biochemical strategies impart bioactivity to devices based on bioinert materials (e.g., silicon^19^, Parylene C, gold^20^, platinum^21^) by presenting favorable biochemical cues. Natural and synthetic bioactive molecules (e.g., growth factors, basement membrane proteins, synthetic peptides and small-molecule drugs^22^) have been incorporated through physical adsorption or chemical immobilization^22,23^. These bioactive modifications have been shown to reduce inflammation and scarring^24,25^ as well as improve recording quality^19^.

Integrating these above principles, devices that are both soft *and* bioactive have the potential to further enhance biointegration by better mimicking the native cell milieu. To achieve this, a strategy proven in tissue engineering to facilitate cell attachment, promote native-like phenotypes, and enhance tissue repair^26,22,23^ is to harness biomaterials inspired by or derived from the extracellular matrix (ECM). As the primary structural and biochemical component of the cellular microenvironment, the ECM provides both mechanical cues and conserved signaling motifs that synergistically regulate cell behaviors^27^. A range of proteins (e.g., silk), polysaccharides (e.g., agarose and chitosan), and synthetic materials have been engineered with bioactive motifs to mimic ECM structure and mechanics. Yet mammalian ECM, such as collagen, elastin, and hyaluronic acid, is the most bioactive for integration in human healthcare applications^28,29^.

Incorporation of ECM-related materials in bioelectronics has often relied on surface modification strategies. Bioactive molecular coatings are insufficient to create a soft microenvironment for surrounding cells, which sense mechanical properties on the micrometer scale^30^. Further, these nanoscale coatings of bioactive molecules are prone to rapid degradation^19^ and thereby, loss of bioactivity. Alternatively, ECM materials of micrometer-scale thickness have been presented as encapsulation layers^20,31^ of bioelectronic devices, which we define as the outermost device layers that act as a physical barrier and structural support by enveloping the conducting material. Of prior work presenting ECM encapsulation layers, silk,^18,32^ albumin^33^, and gelatin^34^ have been investigated with success. In general, reports on non-mammalian ECM encapsulation layers far outweigh those on mammalian ECMs, despite their superior bioactivity and potential for clinical translation. One under-investigated material is collagen. As the most abundant native ECM component in mammals, collagen intrinsically supports biological integration by facilitating tissue remodeling at implantation sites^26,35^, enhancing host cell infiltration^36^, and directing stem cell differentiation *in vitro*^37^. Specifically, type I collagen presents a uniquely promising material for bioelectronic encapsulation. The fibrous structure of type I collagen allows it to be processed into robust, free-standing films that can serve as mechanically stable device carriers^38^. Despite these favorable attributes, collagen has seldom been adopted as an encapsulation material for bioelectronics^31,20,38^, primarily due to its incompatibility with conventional conductors and microfabrication processes. Firstly, the hydrophilicity of collagen molecules promotes water absorption and swelling under physiological conditions. When combined with non-swelling conductors such as metals or metal oxides, this mismatch can induce localized stress at the interface, leading to cracking or delamination of the conducting layer^31^. Secondly, collagen is sensitive to conditions that are utilized in traditional microfabrication processes. The denaturation temperature of fibrous collagen is ∼60–65 °C^39^, and thus requires meticulous control of temperature during thermal evaporation of metallic electrodes^31^. Furthermore, exposure to heat or aggressive solvents^40^ during photoresist development or removal could compromise the integrity of the collagen substrate. Together, these challenges have limited the integration of collagen with traditional electronic materials, leaving its potential as a soft, bioactive encapsulation layer underexplored.

To fully realize collagen’s benefits as bioelectronic encapsulations, there is a need for conducting materials and fabrication strategies that ensure both device stability and collagen bioactivity. Conducting polymers are particularly promising for addressing these requirements. As an alternative to traditional inorganic conductors, conducting polymers are a class of polymers with conjugated backbones that enable electronic conductivity, while their hydrated matrices permit ionic transport, bridging the signaling modalities of electronic devices and biological systems^3^. Their polymeric structures also afford tissue-mimicking mechanical properties^15^, making conducting polymers suitable for integrating with other soft components such as collagen. Among the conducting polymers, poly(3,4-ethylenedioxythiophene):poly(styrene sulfonate) (PEDOT:PSS) is extensively studied due to its commercial availability, solution processibility, tunable mechanical properties^17^, and demonstrated cyto^17,41^- and bio- compatibility^15,42^. Notably, with appropriate additives that promote physical or chemical crosslinking, PEDOT:PSS can form stable, conducting networks at room temperature^43–45^, and could be harnessed to enable integration with temperature-sensitive materials like collagen without compromising biochemical activity.

To build collagen-based bioelectronics with PEDOT:PSS, additive manufacturing, particularly extrusion-based printing, offers distinct advantages. Extrusion-based printing dispenses ink through a nozzle with controlled spatial placement, which can be implemented using readily available syringe-based extrusion and motorized motion systems. Such setups operate under mild, ambient conditions that are inherently compatible with polymer inks and bioactive substrates^46,47^. Owing to its solution processability, PEDOT:PSS has been adopted for extrusion-based printing^12,35,39^, enabling the fabrication of functional bioelectronic devices with diverse and customizable designs. However, the direct printing and stability of conducting polymer electrodes on collagen substrates has yet to be explored.

In this work, we report additive manufacturing strategies for developing collagen-based bioelectronics with PEDOT:PSS electrodes (**Fig. 1a**). We establish extrusion protocols for patterning PEDOT:PSS on collagen films and evaluate their stability, electrochemical performance, and cytocompatibility. We further introduce fabrication strategies for devices fully encapsulated within collagen films. Together, this work presents a versatile platform for fabricating mammalian ECM-based bioelectronics under mild processing conditions. By bridging the gap between soft biological matrices and functional electronic materials, our approach provides a new strategy for developing bioelectronic devices that integrate seamlessly with living systems.

**Figure 1.**
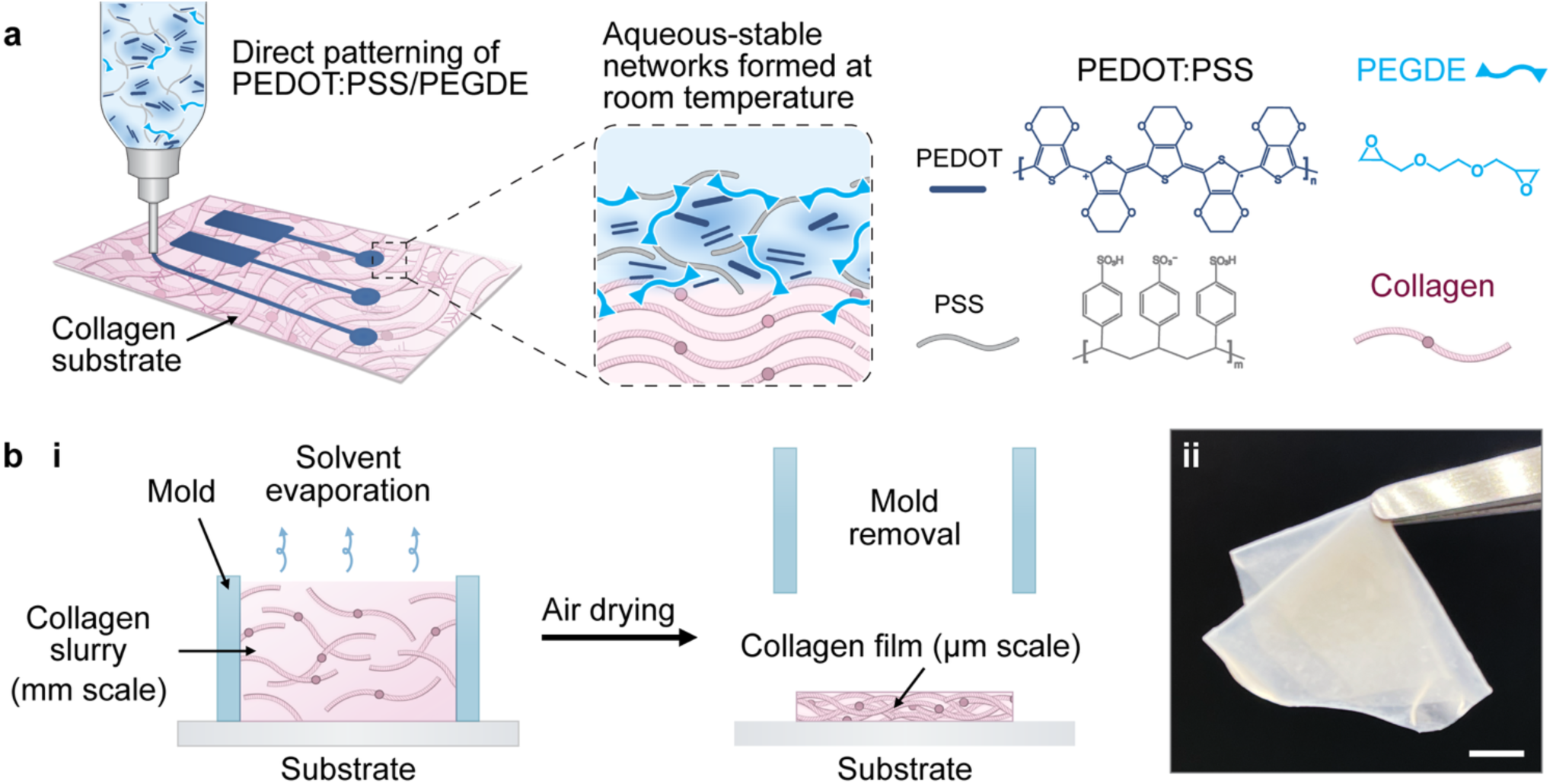
Overview of fabrication and material design for collagen-based bioelectronic interfaces. **(a)** Schematic of direct extrusion printing of PEDOT:PSS/PEGDE ink on collagen substrate at room temperature. PEGDE as a crosslinker has epoxide groups at both ends, which can crosslink with both PSS and collagen. **(b)** Schematic of collagen film fabrication by solvent casting a collagen slurry (mm scale). After air drying, the resulting collagen film (µm scale) can either remain attached to the substrate **(i)** or be handled as a freestanding film **(ii)**. Scale bar = 5 mm.

## 2. Experimental Section

### 2.1. Solvent casting of collagen film

Type I collagen from bovine Achilles tendon (Sigma-Aldrich) was suspended in deionized water at 10 mg mL^-1^ and vortexed until fully wetted. The suspension was then sonicated for 15 minutes and chilled to 4 °C in a refrigerator. Acetic acid (0.05 M) was added, then the mixture was immediately homogenized on medium setting using a VWR 200 homogenizer with a pre-chilled generator probe (10 X 115 mm, saw-tooth) for three 5 second intervals. The collagen slurry was cast at 2 mg collagen per cm^2^ using a chamber slide well over cleaned glass substrate. After air drying in the fume hood, a semi-transparent film was obtained. Collagen film thickness was measured using a micrometer.

### 2.2. Preparation and extrusion 3D printing of PEDOT:PSS/PEGDE ink

PEGDE (M_n_ = 500, Sigma-Aldrich) was added to PEDOT:PSS aqueous dispersion (Clevios PH 1000, 1.0 – 1.3 wt.%) using a positive displacement pipette. The reagents were thoroughly mixed by pipetting up and down followed by vortexing for 1 minute. Finally, the PEDOT:PSS/PEGDE mixture was filtered through a 5 µm polytetrafluoroethylene syringe filter (Cytiva Whatman™). All samples were extruded using a R-GEN 100 3D Bioprinter (RegenHu) equipped with a Volumetric Strand Dispenser printhead (RegenHu). The PEDOT:PSS/PEGDE inks were loaded into a customized Hamilton syringe (RegenHu) and printed through a 30-gauge needle (inner diameter=159 μm, length = 25.4 mm, SAI infusion technologies) with a printhead moving rate of 3 mm s^-1^. All printing patterns were designed and generated in SHAPER (RegenHu). After printing, all samples were dried overnight in ambient conditions and subsequently washed in deionized water overnight.

### 2.3. Tube inversion and rheological characterization of PEDOT:PSS/PEGDE ink

To assess PEDOT:PSS/PEGDE ink stability, 2 mL of freshly prepared ink was transferred into 4.5 mL glass vials. At designated time points (up to 3 days), vials were inverted to evaluate gelation. Samples that flowed to the bottom were classified as liquids, whereas those that remained in place were classified as gels.

Rheological properties of PEDOT:PSS/PEGDE ink were characterized using a TA Instruments HR-20 rheometer with a cone-plate geometry (stainless steel, 40 mm diameter and 1.98889° cone geometry) and a solvent trap to minimize evaporation. Viscosity was measured via rotational flow sweeps at 25 °C across a shear rate range of 0.1-1000 s^−1^. To evaluate stability of the PEDOT:PSS/PEGDE inks over time, oscillatory time sweeps were performed at 25 °C, 1% strain, and an angular frequency 10 rad s^−1^ for up to 5 hours. Ink age was defined as the elapsed time following filtration (∼3 minutes after PEGDE addition). Representative data from multiple independent measurements (N = 3–5) are presented.

### 2.4. Aqueous stability and conductivity measurements

Initial evaluation of aqueous stability was conducted by submerging PEDOT:PSS/PEGDE prints on collagen in deionized water. For qualitative evaluation, images were taken with a stereoscope (Nikon SMZ1270) before and after water addition. Visible dissolution or delamination of the prints were determined as unstable.

For conductivity measurements, PEDOT:PSS/PEGDE samples were prepared by extrusion printing (7 X 7 mm, 90% infill, 2 passes) onto glass or collagen substrates. All samples were measured while hydrated with deionized water. Sample dimensions were measured to calculate conductivity. The thickness of the PEDOT:PSS/PEGDE layer was determined using a micrometer by subtracting the average thickness of the collagen substrate from the total sample thickness (PEDOT:PSS/PEGDE + collagen). Length and width of the conducting samples were obtained from light microscopy images captured with a stereoscope (Nikon SMZ1270) and analyzed using NIS-Elements (Nikon). Conductivity was measured using a four-point probe with soft-tipped probe head (Ossila) under a 0.1 V voltage limit to prevent conducting polymer degradation. For each sample, 50 readings were collected and averaged using the Ossila Sheet Resistance software. Each sample was measured 2 to 3 times, and mean conductivity values were reported.

To evaluate the effects of cell culture conditions on PEDOT:PSS/PEGDE conductivity, samples were first disinfected overnight in 70% ethanol, then transferred to either PBS (1X) or Dulbecco’s Modified Eagle Medium (DMEM, Corning) with 1% penicillin–streptomycin (1%, 10,000 U mL^−1^ penicillin and 10,000 μg mL^−1^ streptomycin, Gibco). Samples were incubated at 37 °C in a cell culture incubator with 5% CO₂ and 95% relative humidity. Prior to conductivity measurements after these incubations, samples were washed 3 times with deionized water (15 minutes per wash) to remove absorbed media.

### 2.5. Stiffness measurements with nanoindentation

The stiffness of hydrated collagen and PEDOT:PSS/PEGDE samples were determined using a Piuma nanoindenter (Optics11 Life) with a spherical-tip cantilever (radius = 28 µm, cantilever stiffness = 206.9 N m⁻¹).

All samples were submerged and equilibrated in PBS (1X), then measured at room temperature. Prior to testing, the cantilever bending was calibrated in PBS on a glass substrate. For each sample, 9 indentation points were collected with a 200 µm spacing between individual indentations. The resulting force–indentation curves were analyzed using the DataViewer software (Optics11 Life) and fitted with a Hertzian contact model to determine the effective Young’s modulus.

### 2.6. Preparation for cell culture and cell seeding

To visualize cell behaviors on different materials, lines of PEDOT:PSS/PEGDE were printed on collagen substrates following the same printing parameters described above. Samples were disinfected in 70% ethanol overnight and washed thoroughly with sterile PBS to remove residual ethanol. Following our previously established protocol, samples were preconditioned by incubating in fetal bovine serum (FBS) overnight in a cell culture incubator^17^. Human umbilical vein endothelial cells (HUVECs, Lonza CC-2519) were maintained following the manufacturer’s protocol and utilized at passage 4 to 5. Prior to cell seeding, preconditioned samples were washed in PBS for 15 minutes to remove excess serum, then incubated in endothelial growth medium (EGM® Endothelial Cell Growth Medium BulletKit®, Lonza CC-3124) for at least 30 min. HUVECs were harvested using ReagentPack™ Subculture Reagents (Lonza CC-5034). The cell suspension (47,000 cells, 1 mL) was seeded dropwise onto each sample at a target density of 5,000 cells cm⁻². Culture medium was changed every other day.

### 2.7. Analysis of cell-material interactions

To determine cell attachment and viability, seeded samples were stained with the LIVE/DEAD kit (Invitrogen, 2 μM Calcein AM, 4 μM of ethidium homodimer-1) in PBS. Fluorescence imaging was performed using a Nikon Eclipse Ts2-FL inverted microscope. Due to the opacity of the collagen film, samples were inverted during imaging to allow visualization of cells on the surface. The total number of cells was determined as the sum of the green-stained live cells (Calcein AM) and red-stained nuclei of dead cells (ethidium homodimer-1). Cell viability (%) was calculated as the proportion of live cells relative to the total cell count.

For cell proliferation analysis, all samples were transferred to microtubes and stored at −20 °C until analysis. To quantify DNA content, samples were lyophilized, digested in Proteinase K solution (100 µg mL^-1^ in 5 mM Tris-HCl 1mM CaCl_2_ buffer, pH 8.0) overnight at 60 °C, and analyzed using the Quant-iT PicoGreen dsDNA assay (Invitrogen) following the manufacturer’s protocol. Fluorescence intensity of cell-free controls was subtracted from experimental groups. DNA quantification was converted to cell number by analyzing cell-seeding aliquots of a known cell number (counted by hemocytometer).

To visualize cell morphology, samples were fixed with 2.5% glutaraldehyde and 2% paraformaldehyde in PBS for 1 hour at room temperature, followed by overnight fixation at 4 °C. Fixed samples were washed and stored in PBS until staining. Cell membranes were labeled with 5 μg mL^−1^ wheat germ agglutinin conjugated with CF®488A (biotium), and nuclei were stained with 10 μg mL^−1^ Hoechst 33342 (BD). After 30 minutes of staining, samples were washed 3 times with PBS prior to imaging.

### 2.8. Electrode device fabrication

Devices for *in vitro* measurements were fabricated by first patterning gold counter electrodes (40 × 20 mm) on glass substrates, briefly described as follows. Adhesive-backed Mylar sheet (Stencil Ease, 5 mil) was laser cut to create a negative mask and placed on cleaned glass slides (25 × 75 mm). Through the mask, a 10 nm chromium layer followed by 100 nm gold was deposited via thermal evaporation (Edwards 306 Vacuum Coater). After the Mylar stencil was removed, the slides were cleaned and cast with collagen films on the opposite end using a modified two-well chamber slide (Nest). PEDOT:PSS/PEGDE working electrodes (2 mm diameter) were then printed on the collagen substrate. To form a sealed chamber for electrolyte, a one-well chamber (Nest) was attached using polydimethylsiloxane (Krayden Dow Sylgard 184 kit) and mechanically secured with the chamber frame and gasket (Nest). For electrical connection, wires were attached to PEDOT:PSS/PEGDE pads using silver-filled epoxy (AI Technology) and secured with silicone sealant (DOWSIL™ 734).

### 2.9. Electrochemical characterization of PEDOT:PSS/PEGDE electrodes

Electrochemical performance of the *in vitro* device was evaluated using a potentiostat (PGSTAT128N, Metrohm). Measurements were programmed in NOVA software (Metrohm) and performed in 5 mL of electrolyte (PBS or DMEM) at room temperature. Electrochemical impedance spectroscopy (EIS) measurements were conducted with a two-electrode configuration (working electrode: PEDOT:PSS/PEGDE; counter electrode: gold) at 0V DC bias versus the open-circuit potential. 0.01 V AC voltage was applied (10^-1^ Hz to 10^5^ Hz).

For cyclic voltammetry (CV) measurements, a three-electrode configuration was used with an additional Ag/AgCl reference electrode (Metrohm, 6.0726.100). The PEDOT:PSS/PEGDE working electrodes were first pre-conditioned by running CV for 20 cycles (−'1 to 0.4 V, 0.02 V/s) to ensure consistent measurements. Immediately afterward, CV measurements were performed for 5 cycles (−0.8 to 0.4 V, 0.1 V/s). Charge storage capacity (CSC) was calculated from the integrated area under the CV curve using a MATLAB script according to the following equation:

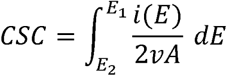

where *E_2_□−□E_1_*is the potential range, *i* is the current at each potential, *v* is the scan rate, and *A* is the area of the PEDOT:PSS/PEGDE working electrode.

Chronoamperometry was performed using a three-electrode setup. Biphasic pulses at 1 s (±0.5□V vs. Ag/AgCl) were applied. The charge injection capacity (CIC) was calculated from the measured current and voltage using:

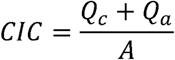

where *Q*_c_ and *Q*_a_ are the total charges injected during the cathodal and anodal phases, respectively, and *A* is the area of the PEDOT:PSS/PEGDE electrode.

To assess the stability of electrode electrochemical performance under cell culture conditions, devices were disinfected overnight in 70% ethanol and washed with PBS. The devices were then incubated in either PBS or DMEM with 1% Pen/Strep in a humidified cell culture incubator. Prior to all measurements, devices were equilibrated at room temperature for 25 min, and temperature stability was verified using an infrared thermometer.

### 2.10. Rheological characterization of cacao butter

For rheological characterization, cacao butter wafers (Navitas Organics) were manually fragmented into ∼5 mm pieces using a spatula and prewarmed in a capped 12 mL syringe submerged in a water bath (∼30 °C) to facilitate loading. The warmed cacao butter was then extruded onto a Peltier plate set to 33 °C, and measurements were performed using a 40 mm cross-hatched parallel-plate geometry (1 mm gap). Prior to testing, samples were equilibrated at 33 °C until viscosity reached a steady state, ensuring a uniform starting condition. All oscillatory measurements were conducted at an angular frequency of 10 rad s⁻¹. Temperature ramp experiments were performed from 20 to 40 °C at a rate of 0.5 °C min⁻¹. To characterize cooling-induced solidification, oscillatory rheology was performed on prewarmed cacao butter at 33 °C while the temperature was rapidly switched to 20 °C. Viscosity and shear-thinning behavior were evaluated using rotational rheology at 33 °C with shear rates ranging from 0.1 to 1000 s⁻¹.

### 2.11. Patterning of collagen encapsulation using cacao butter

To spatially pattern collagen encapsulation, cacao butter was deposited as a sacrificial material directly onto PEDOT:PSS/PEGDE electrodes printed on collagen substrates using a pneumatic melt dispenser (PMD, RegenHu) equipped with a 21G conical dispensing tip (Micron-S, Fisnar). Cacao butter was loaded into a stainless-steel cartridge (RegenHu) and equilibrated at 33 °C for at least 30 min prior to printing. Sacrificial features were aligned to the underlying electrode geometry using SHAPER software (RegenHu). After printing, the cacao butter structures were allowed to solidify at room temperature. Collagen slurry (10 mg mL^-1^), prepared as described above, was then cast over the patterned samples at 2 mg collagen per cm^2^ to form the encapsulation layer. Following overnight drying, cacao butter was removed by immersion in warm deionized water (36 – 37 °C). Samples were gently rinsed by repeated pipetting until no visible oil droplets remained at the water surface. Electrode impedance was measured before and after encapsulation patterning as described above.

## 3. Results and Discussion

### 3.1 Extrusion printed PEDOT:PSS with PEGDE crosslinker forms aqueous stable and conducting patterns on soft, collagen substrates

To fabricate collagen-based bioelectronics with 3D printed PEDOT:PSS electrodes, our objective here was to develop a PEDOT:PSS ink that can firstly extrude and be patterned, yet afterwards maintains an aqueous stable network for biointerfacing and electronic operation. Transitioning from commercially available PEDOT:PSS dispersion to a stable, conducting hydrogel typically requires the incorporation of network-inducing additives and elevated temperatures (60 - 130 °C)^15,49,41,17^ to promote phase separation of PEDOT and PSS domains and thus enable π–π stacking between PEDOT chains^15,17,50^. In some cases, these additives also induce chemical crosslinking of PSS^51^. However, high-temperature processing is incompatible with collagen, which denatures above ∼60 °C^39^. To enable network formation under mild conditions, several chemicals have been shown to induce physical or chemical crosslinking of PEDOT:PSS at room temperature, including dodecylbenzenesulfonic acid^45^, divinylsulfone^43^, and poly(ethylene glycol) diglycidyl ether (PEGDE)^44^. Among these crosslinkers, PEGDE is particularly attractive due to its reactivity to a variety of functional groups^44,52^. The epoxide end groups of PEGDE can react with sulfonate groups in PSS^44^ as well as hydroxyl, amine, and carboxyl groups in collagen^53^. Here, we sought to investigate if PEGDE could be a suitable additive to PEDOT:PSS inks for 3D printing that would not only enable an aqueous-stable, crosslinked PEDOT:PSS network, but also facilitate adhesion of the PEDOT:PSS to the underlying collagen substrate (**Fig.1a**).

First, we examined the compatibility of PEDOT:PSS/PEGDE mixtures for extrusion 3D printing, where compatibility is defined as maintaining a liquid or weak gel state that facilitates extrusion through a nozzle^17,47^. Otherwise, significant gelation during printing can cause excessive extrusion pressure, nozzle clogging, and shearing of the partially formed conducting matrix, which results in fragmented paths that compromise conductivity^17,41^.

PEDOT:PSS/PEGDE mixtures have been reported to form aqueous stable networks without elevated temperature^44^, albeit with techniques that induce solvent evaporation, including drop casting, spin coating, and lyophilization^44,52^. Therefore, we first examined the gelation behaviors of PEDOT:PSS/PEGDE mixtures at room temperature (20 - 25°C) in closed vials, as would be used in 3D printing, to investigate potential gelation in absence of significant solvent evaporation. Colloidal dispersions of PEDOT:PSS were mixed with different concentrations of PEGDE (0, 1, 3, 5, 10 and 15% v/v) and were tested via tube inversion. None of the tested formulations solidified during 3 days of testing, as indicated by flow of the mixture upon vial inversion (**Fig. 2a, Fig. S1**). Consistent with these qualitative observations, rheology of a given formulation (PEDOT:PSS with 5% v/v PEGDE) confirmed that the mixture remained a liquid (loss modulus (G”) > storage modulus (G’), δ > 60°, tan(δ) > 1) for at least 5 hours at room temperature (**Fig. 2b**). Next, to further evaluate PEDOT:PSS/PEGDE as a potential ink, we measured its viscosity across a range of shear rates. The PEDOT:PSS/PEGDE (5% v/v) ink showed a viscosity of ∼0.03 Pa·s at a shear rate of 0.1 s^-1^ (**Fig. 2c**), within the reported range for inks that support smooth extrusion through a nozzle (10⁻²–10LJ Pa·s at 0.1 s⁻¹)^54^. The formulation also exhibited moderate shear-thinning behavior typical of low viscosity inks^55^, with viscosity decreasing as shear rate increased (>50% decrease from 0.1 s⁻¹ to 1000 s⁻¹). Overall, the rheological properties of PEDOT:PSS/PEGDE suggest that it could serve as an ink for extrusion printing.

**Figure 2.**
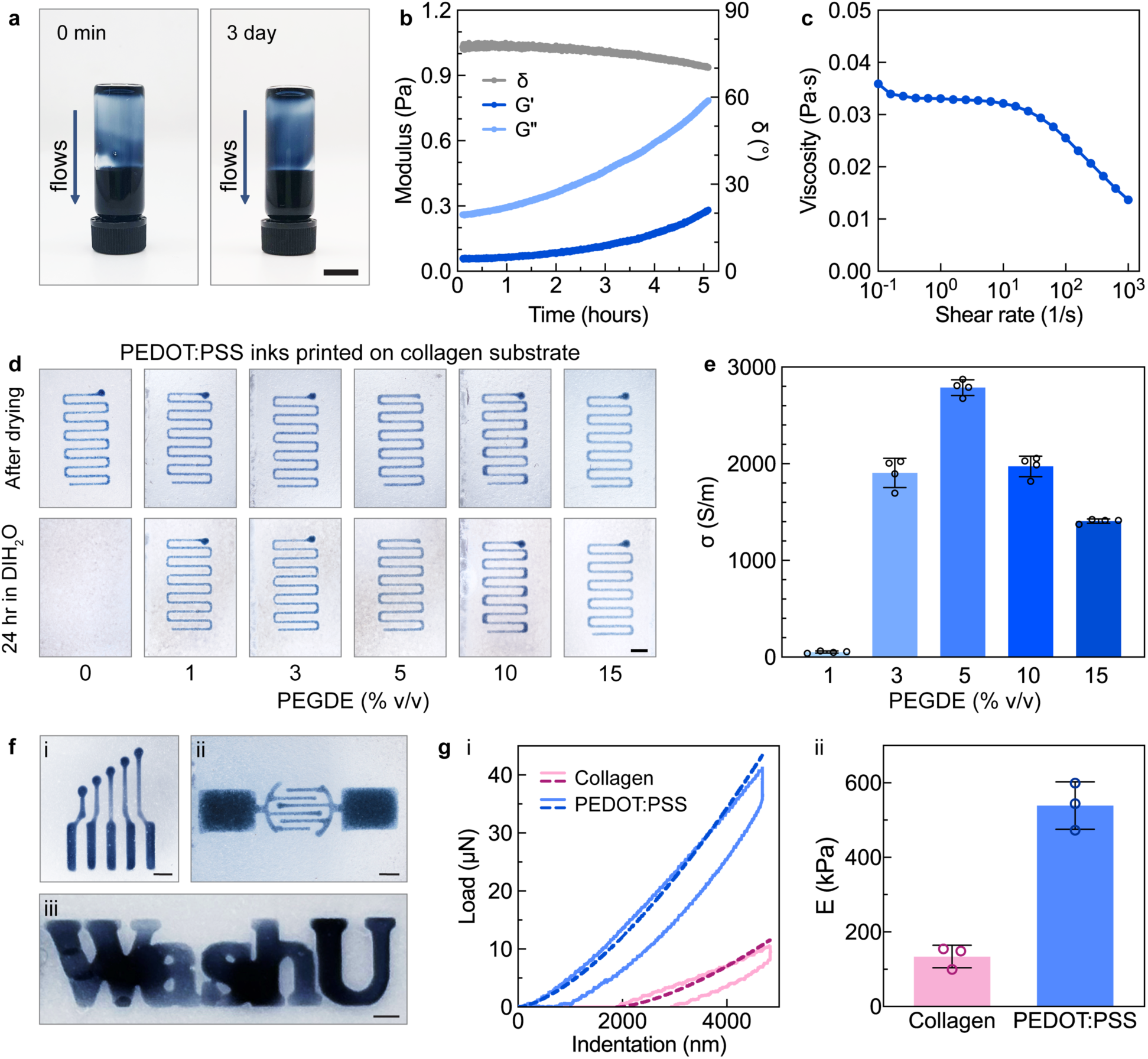
Extrusion printed PEDOT:PSS with PEGDE crosslinker forms aqueous stable and conducting patterns on soft, collagen substrates. **(a)** PEDOT:PSS with PEGDE remained a liquid at room temperature after 3 days, as determined by tube inversion. PEDOT:PSS w/ 5% v/v PEGDE ink shown here as a representative formulation. Scale bar = 1 cm. **(b)** Representative time sweep of storage modulus of PEDOT:PSS with 5% v/v PEGDE (1% strain, 10 rads s^-1^). The mixture remains a liquid after 5 hours at room temperature (loss modulus (G”) > storage modulus (G’), δ > 60 °). **(c)** Viscosity of PEDOT:PSS with 5% v/v PEGDE modestly decreases with increasing shear rate. Representative flow sweep shown. **(d)** PEGDE (1% v/v and above) facilitated formation of aqueous-stable PEDOT:PSS networks on collagen substrates. PEDOT:PSS with 0-15% v/v PEGDE were extruded onto collagen substrates and dried in ambient conditions (top). Without PEGDE, water addition resulted in dispersion of patterned PEDOT:PSS, whereas PEGDE-crosslinked prints remained intact after 24 hours in deionized water (bottom). Representative images, scale bar = 2 mm. **(e)** Conductivities of PEDOT:PSS with different concentrations of PEGDE printed on collagen were compared when hydrated with deionized water. Conductivity increased with PEGDE content up to 5% v/v, then decreased at higher concentrations. N = 4. **(f)** Images of multi-channel electrodes array **(i)**, interdigitated electrodes **(ii)** and the letters “WashU” printed on collagen. Samples printed via a 30G needle with the minimum feature size of 340 μm. Scale bars = 2 mm. **(g)** Representative nanoindentation force-displacement curves **(i)** and Young’s modulus values **(ii)** for collagen (134 ± 30.2 kPa) and PEDOT:PSS/PEGDE (5% v/v) patterned on collagen (539 ± 63.4 kPa). N = 3. Solid lines represent indentation data and dashed lines indicate Hertzian model fits. For **e** and **g**, mean and standard deviation presented. Plot points denote individual samples.

Next, we tested the printing of PEDOT:PSS/PEGDE inks with the same varying PEGDE concentrations on collagen substrates. Collagen substrates were prepared by solvent casting and air drying homogenized type I collagen suspension with acetic acid (**Fig. 1b**). Firstly, all tested PEDOT:PSS-PEGDE mixtures readily extruded through a 30-gauge needle (inner diameter=159 μm). While these mixtures spread when extruded onto glass, these inks maintained their printed pattern on collagen substrates (**Video S1**). The shape retention of liquid PEDOT:PSS/PEGDE was likely due to differences in printed surface roughness, as glass is smooth while fibrous collagen films present a rougher surface^56^. Next, the prints were dried overnight under ambient conditions, as others have used solvent evaporation to induce network formation of these mixtures^44^. To evaluate aqueous stability, defined by resistance to dispersion and stable adhesion to collagen, the dried prints were exposed to deionized water. When introduced to deionized water, PEDOT:PSS prints without PEGDE crosslinker at first partially delaminated (within 3 seconds), then completely disintegrated (within 20 seconds, **Fig. S2**). In contrast, with PEGDE, PEDOT:PSS networks stayed adhered on collagen without any apparent dissolution or delamination upon immediate addition to deionized water. With further incubation in water, printed PEDOT:PSS with as little as 1% v/v PEGDE was stable for at least 24 hours (**Fig. 2d**). In contrast to collagen substrates, printed features on glass were not stably adhered upon water exposure, indicating absent or weak bonding (**Fig. S3**). This contrast in adhesion between substrate types supports that PEGDE is anchoring PEDOT:PSS to collagen, likely through covalent bonding between epoxide groups and collagen functional groups, while the surface roughness of collagen may further contribute through mechanical interlocking.

To investigate the influence of crosslinking on electronic performance, the conductivity of printed PEDOT:PSS with varying PEGDE concentrations on collagen substrates was evaluated in their hydrated state (**Fig. 2e**). Conductivity increased with increasing PEGDE concentrations until 5% v/v, after which the conductivity decreased. At high concentrations, the conductivity likely decreased due to an increasing fraction of non-conductive polymer within the network^57^. As a comparison, PEDOT:PSS/PEGDE printed on glass exhibited a similar trend (**Fig. S4**). Poly(ethylene glycol) polymers, including PEGDE^44^, have been shown previously to increase PEDOT:PSS film conductivity^57,58^. Increasing the PEGDE concentration from 1 to 3% v/v resulted in a >30-fold increase in conductivity (52 S/m to 1906 S/m). Further increasing the PEGDE content to 5% v/v yielded a conductivity of 2788 S/m (> 50-fold increase compared to samples with 1% v/v PEGDE). This maximal conductivity achieved at 5% v/v PEGDE is comparable to previously reported hydrated PEDOT:PSS fabricated with additive manufacturing (>1000 S/m) ^42,48^. Such materials have been successfully used as bioelectrodes for applications including electrical stimulation for peripheral nerve regeneration^15^ and electrophysiological recording of extracellular action potentials^42^, suggesting that this PEDOT:PSS/PEGDE formulation could be suitable for such applications. Amongst the various formulations tested, there were no apparent differences with respect to printing and aqueous stability, yet 5% v/v PEGDE resulted in the highest conductivity. Therefore, this formulation was selected for all subsequent investigations and should be assumed unless specifically noted otherwise. With this selected 5% v/v PEDOT:PSS/PEGDE ink formulation, various designs were successfully patterned on collagen substrates. Demonstrated patterns include a multi-channel electrode array (**Fig. 2fi**), interdigitated electrodes (**Fig. 2fii**), and arbitrary designs such as WashU lettering (**Fig. 2fiii**), showcasing the versatility of the printing approach.

To determine stiffness of these polymer-based devices, we measured the elastic modulus of regions of only collagen as well as PEDOT:PSS/PEGDE on collagen using nanoindentation with a spherical-tip cantilever. To evaluate the mechanical properties under physiological conditions, samples were swollen and characterized in phosphate-buffered saline (PBS). Under these conditions, the collagen substrate and PEDOT:PSS/PEGDE layers had average thickness of 66.2 ± 1.52 μm and 27.0 ± 2.52 μm, respectively, exceeding the characteristic length scales over which cells sense substrate mechanics (microns to tens of microns^30^). The collagen region had a mean Young’s modulus of 134 ± 30.2 kPa, whereas the patterned PEDOT:PSS/PEGDE regions had a modulus of 539 ± 63.4 kPa (**Fig. 2g**). Our measured values for both materials fall within reported ranges. Printed PEDOT:PSS in thin-film geometries typically spans hundreds of kPa^59^ to low MPa range^42,60^ and hydrated collagen films can range from tens of kPa^61^ to several MPa^62,63^. These devices thus are significantly softer than conventional electronic materials (∼MPa–GPa range) and further, fall within the range of soft biological tissues such as skeletal muscle (5–170 kPa) and skin (35–850 kPa)^64^.

### 3.2 PEDOT:PSS/PEGDE on collagen substrates exhibits stable conductivity in cell culture conditions and supports human primary cells

For bioelectronic applications, materials must maintain both structural integrity and functional electrical properties in biologically relevant environments. Biological fluids such as culture media, blood, and interstitial fluid are rich in ions and biomolecules that can interact with PEDOT:PSS and may alter its polymeric and electrical properties^17,65^. To assess material-level stability, we first evaluated conductivity as a metric for polymer integrity under biologically relevant conditions. Prior to testing, samples were first disinfected by immersing in 70% ethanol overnight. Immediately after disinfection and rehydration, mean conductivity nearly doubled compared to non-treated samples (before ethanol: 2788 S/m; after ethanol: 5257 S/m, **Fig. S5**). This conductivity increase aligns with prior reports that water–polar organic solvent mixtures (e.g., 30% water and 70% ethanol) enhance PEDOT:PSS conductivity by promoting phase separation and polymer rearrangement^66^. When next equilibrated in PBS or Dulbecco’s Modified Eagle Medium (DMEM) for one day in standard cell-culture conditions (37 °C, 95% relative humidity, and 5% CO_2_), conductivities were similar to that before ethanol (3004 S/m after PBS and 2467 S/m after DMEM, measurements performed in deionized water, **Fig. S5**). The reduction in conductivity from exposure to electrolyte solutions buffered at physiological pH (PBS and DMEM) relative to deionized water is consistent with prior literature^40^. To assess longer-term stability, samples were incubated in PBS or DMEM for 28 days in tissue culture conditions. Throughout the 28 days, no visible fragmentation or delamination occurred, indicating structural stability (**Fig. 3a**). Additionally, conductivity remained was consistent over time after incubation in both media and PBS, as supported by a two-way analysis of variance (ANOVA)(effect of time, F = 1.76, P = 0.159, **Fig. 3b**). When analyzing the effect of incubation of media type (PBS or DMEM) on conductivity, the effect was significant with samples incubated in DMEM having a lower conductivity (F = 29.1, P < 0.0001, **Fig. 3b**). This is consistent with our previous findings that PEDOT:PSS hydrogel conductivity decreased after exposure to DMEM, potentially due to absorption of ions and nitrogen-containing compounds such as amino acids in the media^17^. No significant interaction between media type and incubation time was observed (F = 1.05, P = 0.397), indicating that the media-dependent difference remained consistent throughout the study period. Taken together, the changes observed in conductivity due to ethanol disinfection or exposure to buffered saline or media indicate that conducting polymer samples should be equilibrated in their intended media before use to establish an accurate and reliable baseline for device performance.

**Figure 3.**
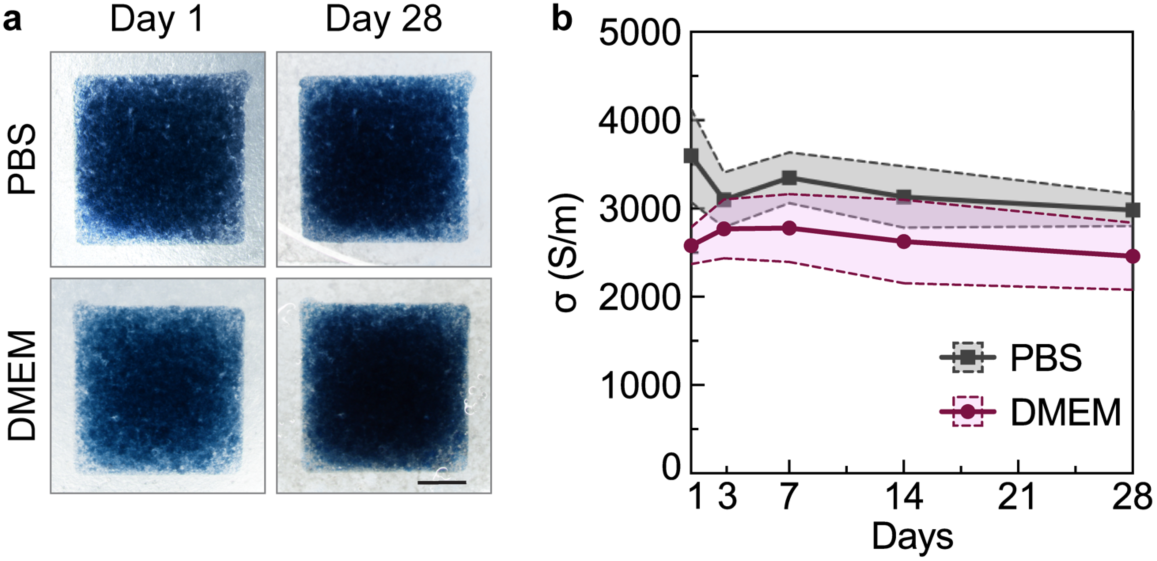
PEDOT:PSS/PEGDE on collagen substrates maintained structural integrity and consistent conductivities in cell culture conditions for at least 28 days. **(a)** Representative images of PEDOT:PSS/PEGDE on collagen samples incubated for 1 day and 28 days in PBS or DMEM. Samples were washed with deionized water before imaging. Scale bar = 2 mm. **(b)** Conductivity of PEDOT:PSS/PEGDE samples in PBS and DMEM remained stable over 28 days of incubation in cell culture conditions (37 °C, 95% relatively humidity, and 5% CO_2_). Two-way analysis of variance (ANOVA) indicated a significant effect of media (PBS vs. DMEM, F = 29.1, P < 0.0001), whereas neither incubation time (F = 1.76, P = 0.159) nor the interaction between media and time (F = 1.05, P = 0.397) was significant. Plot points denote mean and colored shaded areas denote standard deviation. N = 4-5 per group per time point.

Next, we evaluated whether the composition of PEDOT:PSS/PEGDE patterns on collagen could support primary human cell culture. Uniform cell attachment across material interfaces is critical for consistent cell–material interactions, enabling reliable sensing, signal transduction, and tissue integration. While collagen is inherently bioactive, PEDOT:PSS typically requires additional treatment to ensure cell adhesion. Thereby, the substrates were preconditioned with fetal bovine serum (FBS) (**Fig. 4aii**), which we have previously shown enhances early cell attachment^17,41^. This preconditioning enabled uniform seeding of human umbilical vein endothelial cells (HUVECs) across both PEDOT:PSS/PEGDE and collagen regions, resulting in 166% and 173% of the initial seeding density on day 1, respectively (**Fig. 4b**). This increase in cell number after overnight culture indicates support for not only attachment but early proliferation of HUVECs (doubling time 15-48 hours) on both PEDOT:PSS/PEGDE and collagen substrates.

**Figure 4.**
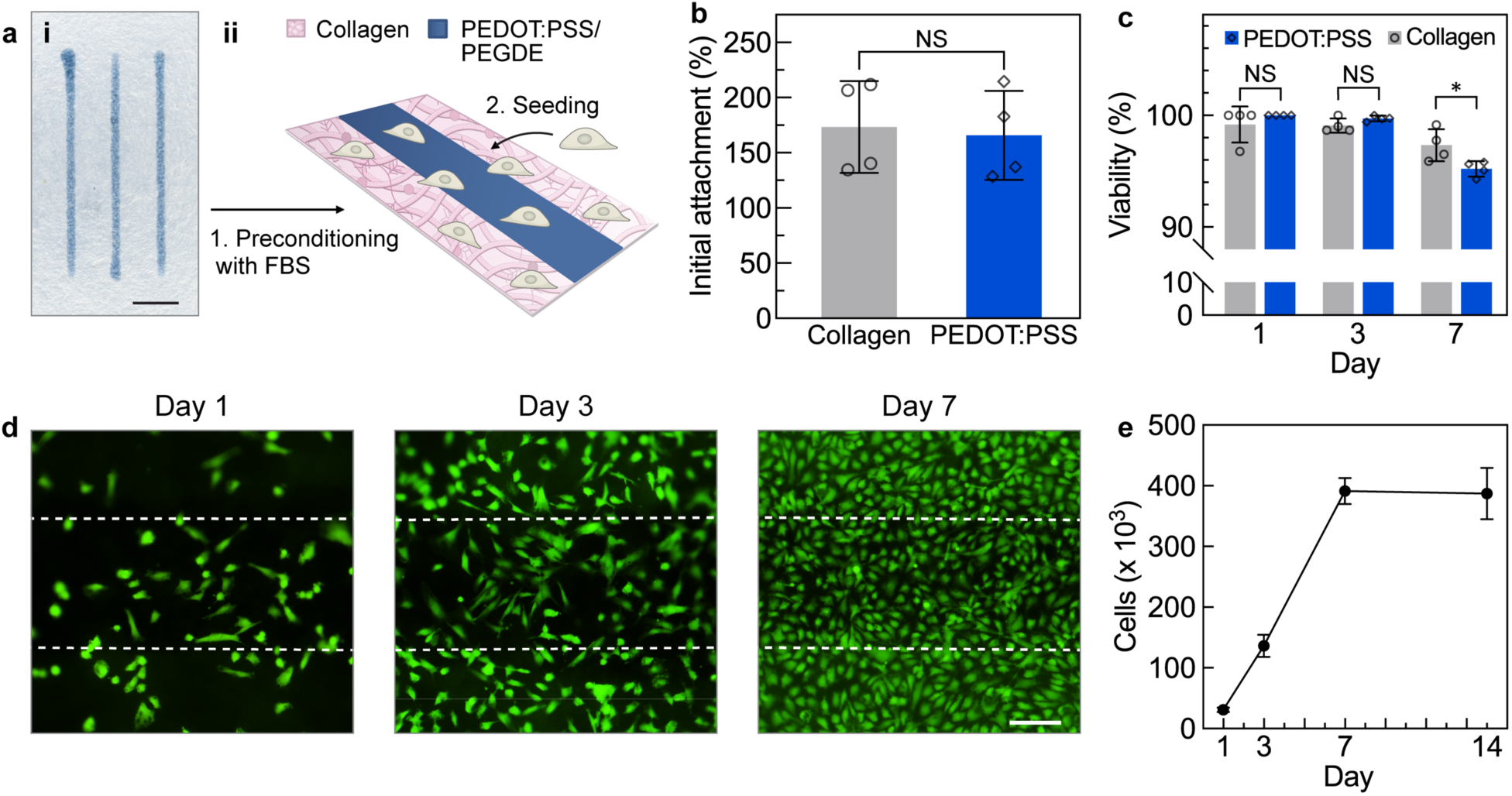
PEDOT:PSS/PEGDE patterned on collagen substrates supports human umbilical vein endothelial cell attachment, proliferation, and formation of a confluent cell layer. **(a)** PEDOT:PSS/PEGDE lines extruded onto collagen for cytocompatibility tests **(i)**. Scale bar = 2 mm. Samples were preconditioned with fetal bovine serum (FBS) **(ii)** prior to seeding human umbilical vein endothelial cells (HUVECs). **(b)** Initial attachment (day 1 cell density as a percentage of seeding density) was high on both collagen and PEDOT:PSS/PEGDE regions. Cell density determined via fluorescence imaging of live cells stained with Calcein AM. N = 4. Unpaired t test revealed no significant effect of cell culture substrates (F = 1.06, P = 0.802). **(c)** Viability of HUVECs on PEDOT:PSS/PEGDE and collagen over 7 days as quantified by Live/dead staining (Calcein AM and ethidium homodimer-1). Two-way analysis of variance (ANOVA) followed by Šidák-adjusted multiple comparison tests, which are shown on the plot. Viability was not significantly affected by substrate type (F = 0.337, P = 0.569), but both culture time (F = 30.0, P < 0.0001) and the substrate–time interaction (F = 5.81, P = 0.0113) contributed significantly. **(d)** Representative images of HUVECs (green, Calcein AM) cultured on PEDOT:PSS/PEGDE on collagen over 7 days. Area bounded by dashed lines indicate PEDOT:PSS/PEGDE tracks. Scale bar = 100 μm. Red channel not shown due to significant auto-fluorescence of collagen. **(e)** Quantification of cell number over time by DNA content. Cell number remained constant from Day 7 to Day 14. N = 4. Mean and standard deviation presented. Plot points denote individual samples. *P ≤ 0.05, non-significant (NS) P > 0.05.

The hybrid substrates robustly supported cell viability, with a viability >95% throughout days 1, 3, and 7 (**Fig. 4c**). Mean viability had a slight decline at day 7, likely associated with the high cell density. Comparison of mean viabilities among collagen and PEDOT:PSS/PEGDE regions were not statistically significant for days 1 and 3. There was a small significant difference at day 7, but overall viability was still high for both groups (P = 0.0179, 95.2% vs 97.3%, **Fig. 4c**). Additional analysis of live cell imaging (Calcein AM, **Fig. 4d**) and DNA quantification (PicoGreen, **Fig. 4e**) both confirmed steady increases in cell number until confluency (day 7). The confluent cell layer was then sustained for an additional 7 days, with a stable cell number from days 7 to 14 (**Fig. 4e**). Morphological analysis further revealed formation of a continuous and physiologically representative endothelial layer. Fixed HUVECs exhibited the characteristic cobblestone morphology spanning both conducting and non-conducting regions (**Fig. S6**). Overall, these results suggest that the device preparation process produces PEDOT:PSS/PEGDE surfaces that support cell attachment and growth comparable to collagen as well as stability in longer-term culture.

### 3.3 PEDOT:PSS/PEGDE electrodes on collagen maintain stable electrochemical performance in cell culture conditions

After establishing the stability of the intrinsic electrical property of conductivity as well as the cytocompatibility of PEDOT:PSS/PEGDE patterned on collagen, we next evaluated the material’s performance when used as electrodes in a tissue-culture environment. Considering that bioelectronic devices must operate in aqueous conditions while maintaining reliable external connections, we designed a containment system to isolate the active area of the device (i.e. electrodes) from device connections (**Fig. 5a**). These contacts supply power and transfer signals, therefore they must remain dry to prevent noise, drift, and unstable operation. We adapted a chamber-slide configuration to have the circular electrode area exposed to cell culture media while minimizing hydration of the interconnects and contacts pads. To fabricate these devices for electrochemical measurements, collagen substrates were cast onto glass slides patterned with gold counter electrodes, followed by extrusion printing of 2 mm diameter PEDOT:PSS/PEGDE working electrodes. Multiple electrodes were printed per substrate to generate technical replicates.

**Figure 5.**
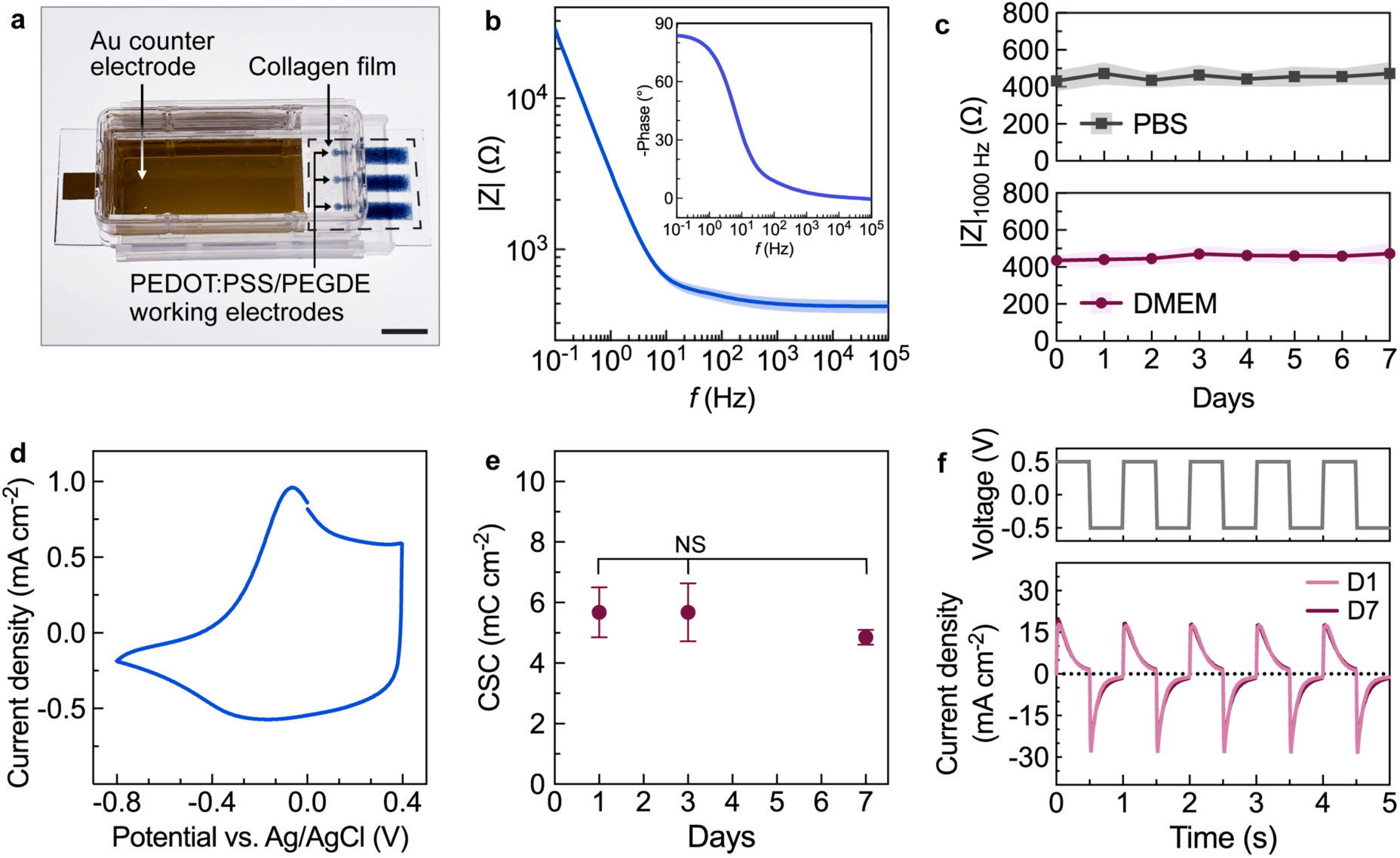
PEDOT:PSS/PEGDE electrodes on collagen maintain stable electrochemical performance in cell culture conditions. **(a)** Photograph of the electrode set-up for electrochemical testing. A chamber slide was modified to contain electrolyte. PEDOT:PSS/PEGDE printed on collagen (bounded by dashed line) served as the working electrodes, with a gold counter electrode completing the circuit. Scale bar = 10 mm. **(b)** Electrochemical impedance spectroscopy measurements in PBS from multiple devices shown as Bode magnitude (|Z| vs f) and the corresponding phase plots (-phase vs f, insert). N = 18. Mean presented as bold line and standard deviation presented as shaded area. **(c)** Electrode impedance at 1000 Hz in PBS and DMEM after incubation (37 °C, 95% relative humidity, and 5% CO^2^). Two-way repeated measures analysis of variance (ANOVA) showed no significant effect of time (F = 1.23, P = 0.289), media (F = 0.0695, P = 0.792) or their interaction (F = 0.418, P = 0.890). Plot points denote mean and colored shaded areas denote standard deviation. N = 9. **(d)** Representative cyclic voltammetry measurement of the device in PBS. N = 3 **(e)** Charge storage capacity (CSC) was not significantly different over 7 days in DMEM (37 °C, 95% relative humidity, and 5% CO^2^). Separate samples were measured at each time point. Mean and standard deviation presented. One-way ANOVA (F = 1.22, P = 0.361) and Tukey’s multiple comparison tests performed. Non-significant (NS) P > 0.05. N = 3. **(f)** Representative biphasic stimulation pulses (top) and the corresponding current density (bottom) for PEDOT:PSS/PEGDE electrode in DMEM on day 1 vs day 7. N = 3. Chronoamperometry performed against an Ag/AgCl reference electrode. Separate samples were measured at each time point.

We first conducted electrochemical impedance spectroscopy on electrodes in PBS. Electrochemical impedance spectroscopy measures the frequency-dependent response of the electrode–electrolyte interface to a small-amplitude AC perturbation. It is widely used to assess neuroelectrode recording performance^67^ as well as to monitor cell behaviors *in vitro* through impedance-based sensing^4,52^. The Bode plots (|Z| and -phase vs frequency) of the electrodes exhibited the characteristic “hockey-stick” profile. At high frequencies, the impedance remained relatively constant with a small phase angle, reflecting predominantly resistive behavior. At lower frequencies, the impedance increased and the phase approached ∼80°, consistent with a capacitance-dominated response. At the conventional benchmarking frequency of 1000 Hz, the electrodes exhibited low mean impedance which was highly consistent across devices (|Z|_1000 Hz_ = 436 ± 43.0 Ω, coefficient of variation = 9.86%). Additionally, the electrodes displayed a low mean cut-off frequency (*f* _cutoff_ = 8.46 ± 1.03 Hz at -phase ∼ 45°), providing a wide operating bandwidth above cutoff where non-linear signal distortion is minimized (**Fig. 5b**).

We next examined stability of impedance in physiologically relevant media (PBS and DMEM). Despite the media-dependent differences in polymer conductivity observed previously, electrolyte composition did not significantly affect impedance as determined by two-way repeated measures ANOVA (|Z|_1000 Hz_ in PBS vs in DMEM, F = 0.0695, P = 0.792). Measured impedance represents a combination of electronic, ionic, and interfacial contributions, hence variations in electronic conductivity alone may have only a minimal overall impact. There was no significant effect of time (F = 1.23, P = 0.289) or interaction between media and time (F = 0.418, P = 0.890) on |Z|_1000 Hz_ over the 7 days of incubation, suggesting impedance stability in cell culture conditions (**Fig. 5c**).

Alternatively to recording, electrodes can be used for electrical stimulation. For stimulation, the electrode needs to drive current over the solid-liquid boundary without changing the electrode properties. To evaluate stimulation-relevant performance, electrodes were subjected to cyclic voltammetry (CV), a standard method for assessing redox stability and charge-storage capacity (CSC). The CV curves displayed the expected parallelogram-like shape characteristic of capacitive PEDOT:PSS behavior (**Fig. 5d**). Additionally, CSC of the electrodes exhibited a linear behavior with scan rate (10 – 500 mV cm^-1^, **Fig. S7**), which is consistent with ideal double-layer capacitive behavior without diffusion limitations for polymer electrodes. The mean CSC value (5.68 ± 0.827 mC cm⁻²) fell within reported ranges for PEDOT-based electrodes^60,68^ and substantially exceeds the threshold of 0.5LJmC cm⁻² required for effective stimulation^68^. The effect of time on CSCs was not significant over seven days when incubated in DMEM (one-way ANOVA, F = 1.22, P = 0.361, **Fig. 5e**) suggesting that the electrode had stable charge storage under these conditions. Additionally, there was no significant effect of time on the peak-to-peak separation values (ΔEp) (**Fig. S8**), defined as the potential difference between the anodic and cathodic peaks in cyclic voltammetry,(one-way ANOVA, F = 2.21, P = 0.191), indicating consistent redox behavior without evidence of progressive interfacial degradation. Next, we quantified charge-injection capacity using biphasic current pulses in cell culture medium (**Fig. 5f**). The electrodes did not have a significantly different charge injection capacity when comparing day 1 to day 7 (unpaired t-test, F = 9.45, P = 0.0577, **Fig. S9**). In conclusion, these characterizations indicate that the fabricated PEDOT:PSS/PEGDE electrodes on collagen could serve as stable recording or stimulating electrodes.

### 3.4 Selective patterning of collagen films using sacrificial cacao butter to create a fully encapsulated device

For bioelectronic devices operating in *in vitro* conditions, a simple electrode-on-substrate configuration can maintain stable operation, as we have demonstrated above. In contrast, implantable bioelectronics are exposed to harsher conditions including surgical handling and tissue dynamics^69,70^, and therefore typically require top and bottom encapsulation layers to prevent mechanical failure (e.g., scratching, cracking, or delamination of the conducting tracks). There have been a few works reporting the use of collagen for device encapsulation, and in these works, the collagen layer was introduced as a continuous film by drop casting^31,20^. Although straightforward, this approach lacks spatial control and fully covers the electrode area. A dense collagen layer, resembling fibrotic capsules formed *in vivo*, is expected to compromise signal quality^70^. Therefore, selective patterning of collagen is necessary to protect device components while keeping the electrode surface exposed. However, precise patterning of collagen films remains challenging. Patterning of conventional electronic encapsulation materials, including elastomers and parylene C, typically rely on methods such as photolithography^15^ and laser micromachining ^69,71^ which are mostly incompatible with collagen.

Alternatively, sacrificial materials have been used to pattern sensitive biomaterials, often using 3D printing. 3D printed sacrificial materials such as carbohydrate glass, gelatin, and Pluronic have been used to create defined channels inside fibrin, Matrigel, and even a bath of ECM matrix and organoids^72,73^. Although compatible with sensitive biomaterials, these strategies are designed for fabricating hydrogels by sol-gel methods. In contrast, these collagen films are made by solvent casting. Solvent-cast collagen begins as an acidic aqueous slurry several millimeters thick^61^, which is subsequently air-dried to form a robust film that is microns-thick (**Fig. 1b**). During this process, the collagen component undergoes extensive change in thickness. Hence, a sacrificial material for this process must (1) be able to support itself in 3D structures, (2) maintain structural integrity in acidic aqueous conditions during solvent evaporation, and (3) be removable under mild conditions expected to preserve collagen’s structure and bioactivity. Existing sacrificial materials including hydrogels^74^, carbohydrate glass^75^ and thermoplastics^76^ fail to meet all of these requirements for solvent casting. To overcome this limitation, we investigated cacao butter as a novel sacrificial material (**Fig. 6a**).

**Figure 6.**
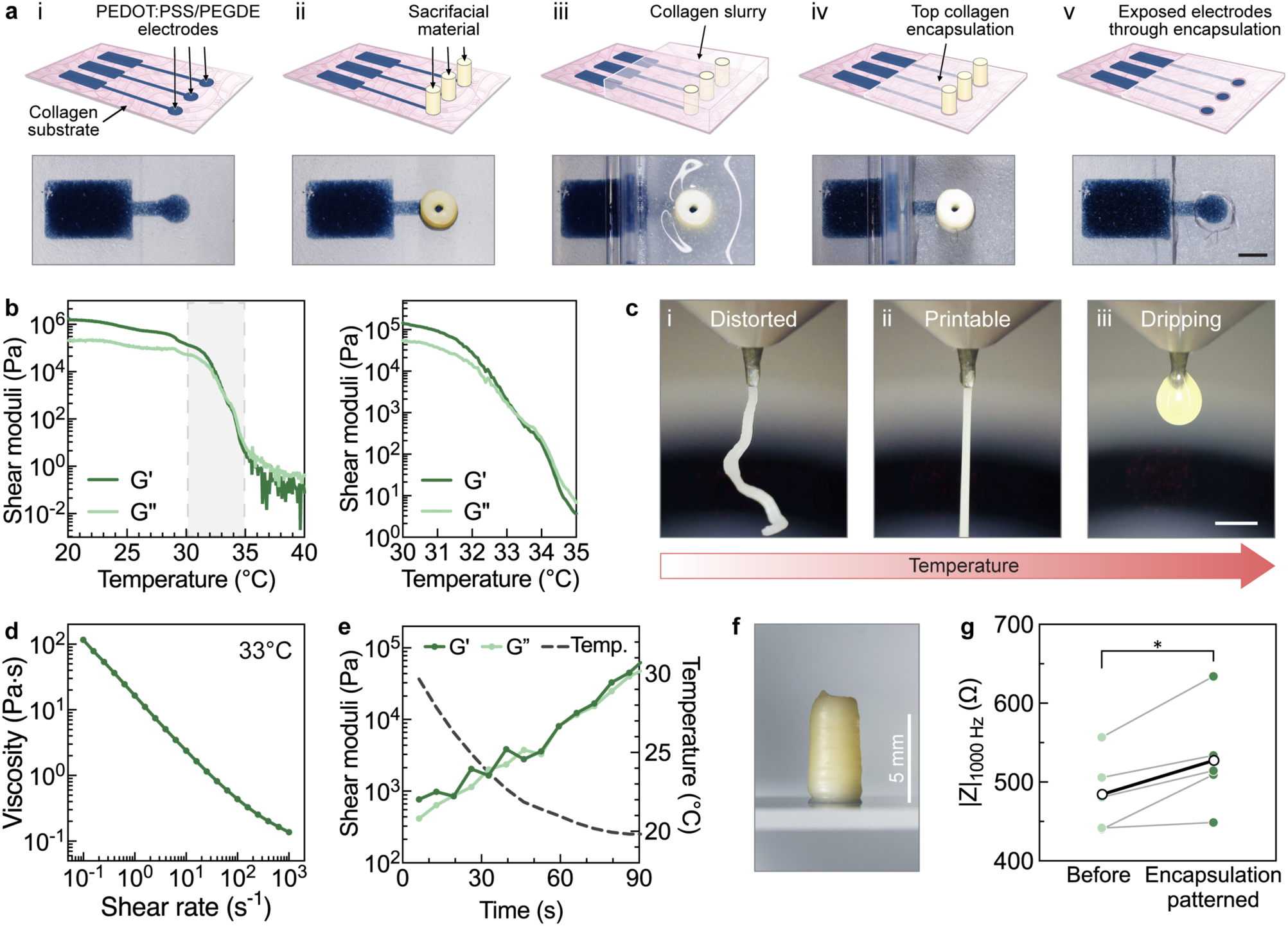
3D printed, sacrificial cacao butter enables selective patterning of collagen encapsulation. **(a)** Schematic (top) and representative photos (bottom) of patterning collagen encapsulation layer with sacrificial cacao butter pillars. Sacrificial pillars were deposited directly onto the PEDOT:PSS/PEGDE electrodes (ii). Next, collagen slurry was cast onto the sample (iii) and air dried to form the collagen film through solvent casting (iv). After removal of the sacrificial pillars by warm water washing, the device was left with exposed electrodes (v). Scale bar = 2 mm. **(b)** Representative temperature ramp of cacao butter shear moduli (0.01% strain, 10 rad s^-1^, 0.5°C/min) from 20 to 40 °C (left), with zoomed in panel of the grey shaded area showing rapid decrease of the storage (G’) and loss (G”) modulus in between 30-35 °C. **(c)** Representative photos of extrusion of cacao butter at 32, 33, and 34 °C through a 23G nozzle (ID=0.564 mm). At 32 °C, the cacao butter required high pressure (100 kPa) to extrude and resulted in a distorted strand **(i)**. When the cartridge temperature was increased to 33 °C, the cacao butter was easily extruded with 20 kPa of air pressure applied. A soft, continuous strand was extruded **(ii)**. With further increased temperature to 34 °C, the cacao butter turned into a liquid. When a smaller pressure was applied (10 kPa), the liquid cacao butter continuously dripped from the nozzle, and thus was deemed unsuitable for printing 3D structures **(iii)**. Scale bar = 2 mm. **(d)** At 33 °C, cacao butter exhibited shear-thinning behavior as its viscosity decreased with increasing shear rate. Representative flow sweep shown. **(e)** After switching from 33 °C to 20 °C, shear modulus of cacao butter rapidly increased (>10 fold within 60 s, 0.01% strain, 10 rad s^-1^), facilitating rentention of the printed shape. Representative run shown. **(f)** Cacao butter could be built into 3D structures that were 5 mm or taller to be suitable for patterning solvent cast films. Representative image, scale bar = 5 mm. **(g)** Impedance of PEDOT:PSS/PEGDE electrodes before versus after patterning top collagen encapsulation. The patterning process resulted in minor increase in impedance of PEDOT:PSS electrodes. Individual measurements (green) and mean values (black and white) presented. N = 4. Repeated measures paired t test (P = 0.0314). *P ≤ 0.05.

Cacao butter is primarily composed of long-chain fatty acids^77^, which impart hydrophobicity that is expected to maintain structural integrity during the aqueous solvent-casting process. Cacao butter also melts around physiological temperatures (35 – 37 °C^77^), which should allow both deposition and removal under conditions that are compatible with collagen. Although chocolate (typically containing 20–55% cacao butter) has been 3D-printed in the food industry, neither cacao butter nor its composites have been reported as sacrificial materials, to our knowledge. To first determine whether cacao butter can be used as a printable material, we conducted qualitative extrusion tests alongside examination of rheological properties to identify material states that could enable smooth, controlled extrusion. We first characterized its shear moduli as a function of temperature from 20 °C (room temperature) to 40 °C (above melting temperature of cacao butter) (**Fig. 6b**). Qualitative evaluation of extrudability was also conducted at selected temperatures within this range using pneumatically driven extrusion 3D printing (**Fig. 6c**). Initially at room temperature, cacao butter behaved like a solid (G’>G”). As the temperature was increased, its storage and loss moduli gradually decreased. Extrusion was attempted at 32 °C at which cacao butter was characterized to be a solid with a high storage modulus (G’>10^5^ Pa). Extrusion was accomplished at high pressure (100 kPa, **Fig. 6ci**); however, the resulting strand was distorted, indicating inconsistent material flow. As the temperature was further increased to 33 °C, both G’ and G” decreased to the 10^3^ Pa range (G’ ∼ 2.11 x 10^3^ Pa, G”∼ 1.73 x 10^3^ Pa, **Fig. 6b**). At this temperature, cacao butter was easily extruded into a continuous filament with 20 kPa of air pressure applied (**Fig. 6cii**). With further elevated temperature, the material became more viscously dominated as evidenced by loss modulus surpassing the storage modulus (G”>G’) (**Fig. 6b**). Cacao butter was no longer able to form continuous strands during extrusion, as observed by the liquefied cacao butter dripping from the nozzle (34 °C, **Fig.6ciii**). Therefore, we identified 33 °C as the optimal printing temperature for the following experiments. We further characterized the viscosity and shear behavior of cacao butter at this identified temperature. At 33 °C, its viscosity decreased with increasing shear rate (**Fig. 6d**), indicating shear-thinning behavior responsible for its ability to extrude.

After extrusion of the cacao butter, the printed construct was maintained at room temperature (∼20 °C) and structures appeared to retain their printed features without collapse. To characterize the rheology of the ink after printing, we switched temperature of the cacao butter from the melting state (33 °C) to room temperature (20 °C) while measuring its properties. A rapid transition to a solid-like behavior (G’>G”) and an increase in the storage modulus of cacao butter by >10 fold was observed within 60 s (**Fig. 6e**). This solidification of cacao butter at room temperature was beneficial for maintaining the printed structure. In our case, we utilized this feature to create structures with millimeter-scale height; the material could support subsequent layers of printing to create 3D structures. By using this layer-by-layer printing, high aspect ratio cacao butter pillars (∼1 mm diameter, >5 mm height; **Fig. 6f, Video S2**) were fabricated to accommodate the initial height needed for solvent casting the collagen encapsulation. The pillars remained intact and adhered to printed PEDOT:PSS/PEGDE electrodes during addition of the aqueous collagen solution (**Fig. 6a.ii-iii**). After overnight drying to produce the collagen film, the cacao butter pillar could be easily removed using warm water of 36 – 37 °C, a temperature that is both near the melting temperature of cacao butter as well as around physiological temperature (**Video S3**). After washing, PEDOT:PSS/PEGDE electrodes were clearly exposed (**Fig. 6a.v**). Impedance measurements of electrodes before patterning cacao butter and after its removal showed a statistically significant yet minor increase of 6.39 % (485 Ω to 516 Ω; repeated measures paired t test, P = 0.0314, **Fig. 6g**), indicating that this process does not substantially affect device performance. In all, we present a novel sacrificial material that may be useful for patterning other biomaterials, especially those made via solvent casting. Specifically for collagen here, we show how cacao butter can be used to fabricate encapsulated microelectrodes.

## 4. Conclusions

In this work, we establish an additive manufacturing approach that integrates PEDOT:PSS electrodes with collagen substrates and encapsulation layers. PEGDE-mediated crosslinking enables aqueous stable, conductive polymer networks to be printed and formed directly on collagen under mild conditions, overcoming previous fabrication incompatibilities. The resulting devices exhibit tissue-like mechanical properties, support cell attachment and proliferation, and maintain consistent electrochemical performance in physiological environments. We further introduce a sacrificial 3D printing strategy using cacao butter to achieve spatial control of collagen encapsulation while preserving electrode function. Together, this work establishes a versatile platform for ECM-based bioelectronics and provides a foundation for developing biointerfaces with improved biological integration for tissue models and implantable systems.

## Author Contributions

T.L. and A.L.R. envisioned the study and wrote this manuscript; T.L. J.P., S.S.O., A.P.G., and A.L.R. were involved in data interpretation; T.L. investigated ink formulations, material stability, and performed all microscopy, electrical and rheological characterization of PEDOT:PSS/PEGDE and cacao butter; J.P. performed nanoindentation measurements; S.K.M. contributed to methodology of sealing well attachment for electrodes and developed MATLAB script to analyze charge storage capacity; B.A.S. and S.S.O. contributed to methodology development and data interpretation for rheological characterizations; R.M.A. troubleshooted melt extrusion; S.K.M, J.S.Y., C.P.O, and Y.W., contributed to materials preparation for fabricating gold electrodes; S.K.M, J.S.Y., C.P.O, Y.W., and C.J.V.E. fabricated collagen films.

## Acknowledgements

This research was supported by the National Science Foundation through GR #0039349 and FR #2319060, and the Center for Engineering MechanoBiology (CEMB), an NSF Science and Technology Center, under grant agreement CMMI #15-48571. The authors also acknowledge support from Washington University in St. Louis through the Women’s Health Technologies Collaboration Initiation Grant, Center for Regenerative Medicine Seed Grant, Ovarian Cancer Research Innovation Fund Award, and the McDonnell Center for Cellular and Molecular Neurobiology Small Grant. S.S.O. acknowledges support from the McDonnell International Scholars Academy. The authors also acknowledge assistance from staff and use of instruments from the Institute of Materials Science and Engineering and Department of Mechanical Engineering and Materials Science. The authors thank Profs. Nathaniel Huebsch and Cory Berkland (Department of Biomedical Engineering, Washington University in St. Louis) for access to an oxygen plasma chamber and plate reader, respectively. The authors acknowledge the use of ChatGPT (accessed April 2026) to assist with reorganization and transitions to improve flow in the Introduction as well as Results and Discussion sections. All suggestions were reviewed, revised, and approved by the authors, who take full responsibility for the accuracy and integrity of the work.

