## Supplemental Information for "Additive manufacturing of PEDOT:PSS electrodes on collagen substrates for soft and bioactive electronics"

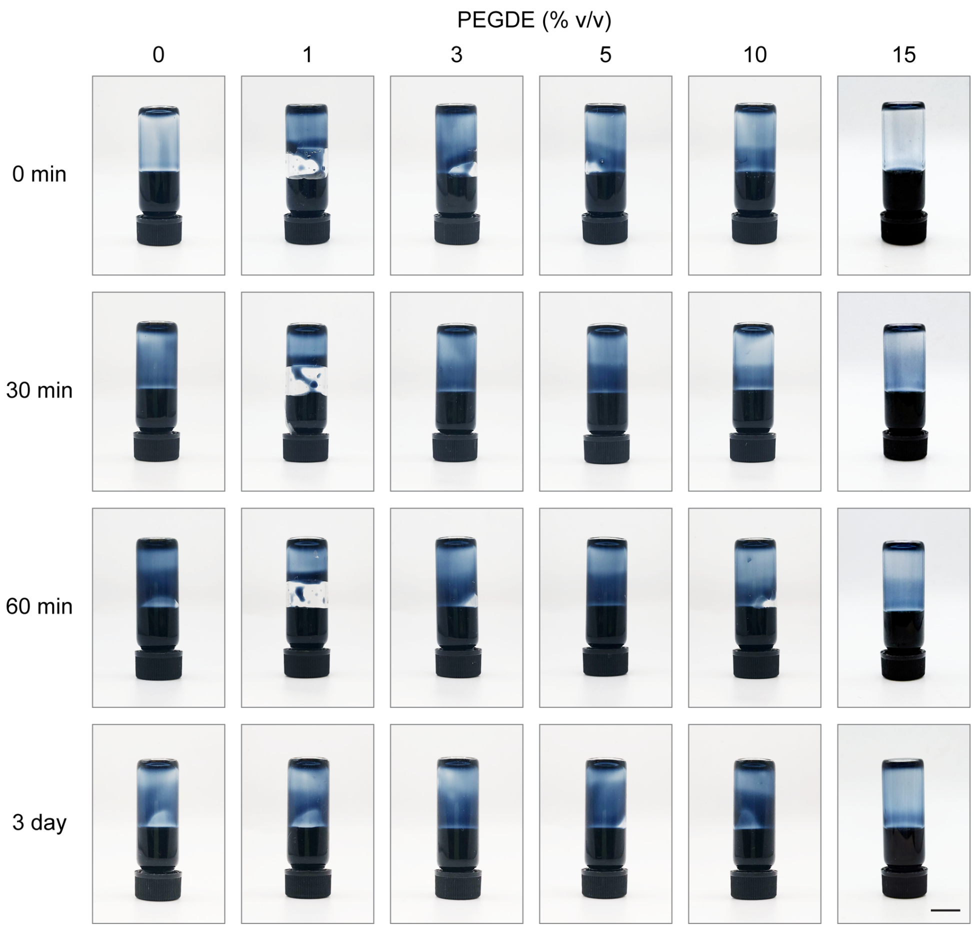


**Supporting Figure 1. PEDOT:PSS with various PEGDE concentrations remained liquid over 3 days, as determined by tube inversion**. The condition of 5% v/v PEGDE at 3 days is repeated here from Figure 1b for the purposes of comparison to other conditions. N=1. Scale bar = 1 cm.

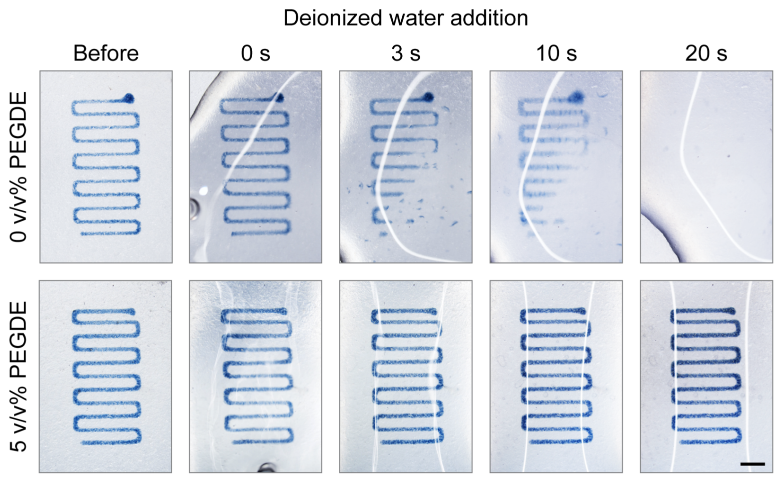


**Supporting Figure 2. PEGDE enhances adhesion and aqueous stability of PEDOT:PSS prints on collagen.** Following exposure to deionized water, PEDOT:PSS prints without PEGDE crosslinker (top) rapidly delaminated within seconds and fully disintegrated within 20 seconds. With the addition of PEGDE (5% v/v), the PEDOT:PSS print remained adhered to the collagen substrate (bottom). Representative images. Scale bar = 2 mm.

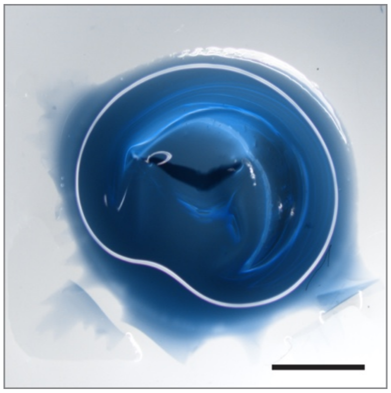

**Supporting Figure 3. PEDOT:PSS/PEGDE deposited on glass delaminated after rehydration.** The film detached from the glass substrate, forming a water-filled gap in the middle, while a small adhered region (lower left corner) resulted in localized tearing. Representative photo shown for PEDOT:PSS with 3% v/v PEGDE. Scale bar = 2 mm.

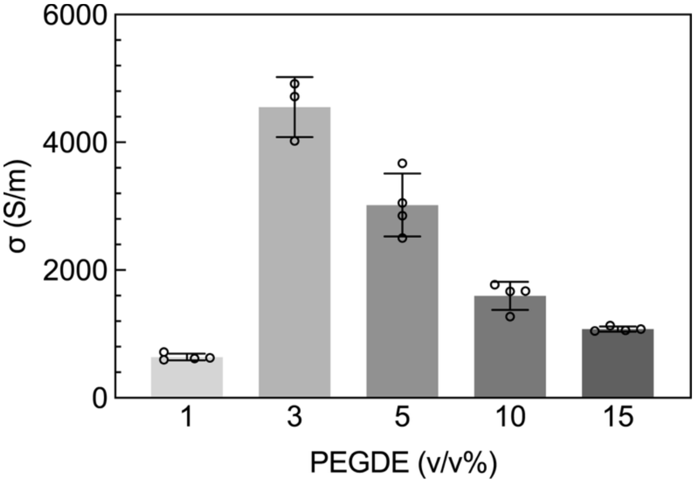


**Supporting Figure 4. Conductivity of PEDOT:PSS with different concentrations of PEGDE printed on glass substrates.** Measurements performed on samples fully hydrated with deionized water. Mean and standard deviation presented. Plot points denote individual samples. N = 3 - 4.

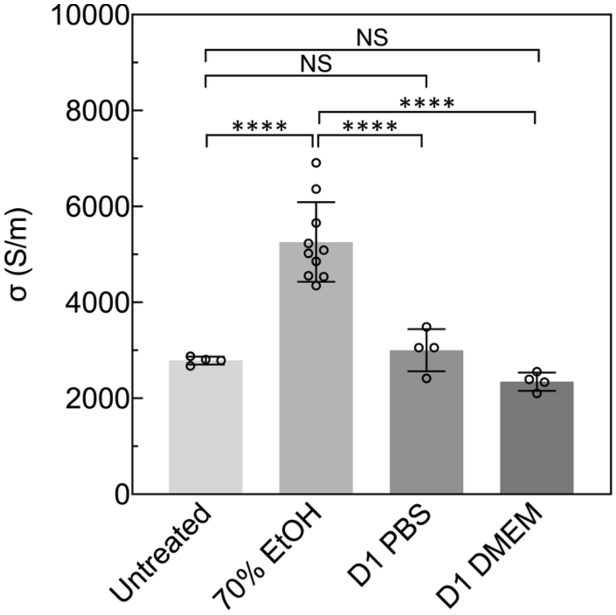


**Supporting Figure 5. Conductivity of PEDOT:PSS/PEGDE on collagen transiently increased after ethanol disinfection, then returned to baseline following 1 day in PBS or DMEM.** Conductivity measurements were performed at various stages of sample preparation for cell culture. Upon disinfection with 70% ethanol, conductivity increased; but after phosphate buffer saline (PBS) or Dulbecco's Modified Eagle Medium (DMEM) immersion for 1 day, conductivity was similar to that of undisinfected samples. All samples were thoroughly washed in deionized water prior to conductivity measurements. N = 4-10. The untreated PEDOT:PSS/PEGDE (5% v/v) condition is repeated from Figure 1f, and the 1-day PBS and DMEM incubation conditions are repeated from Figure 2b for comparison. Mean and standard deviation presented. Plot points denote individual samples. One-way analysis of variance ($F = 31.4$, $P < 0.0001$) and Tukey's multiple comparison tests performed. ****$P \leq 0.0001$ non-significant (NS) $P > 0.05$.

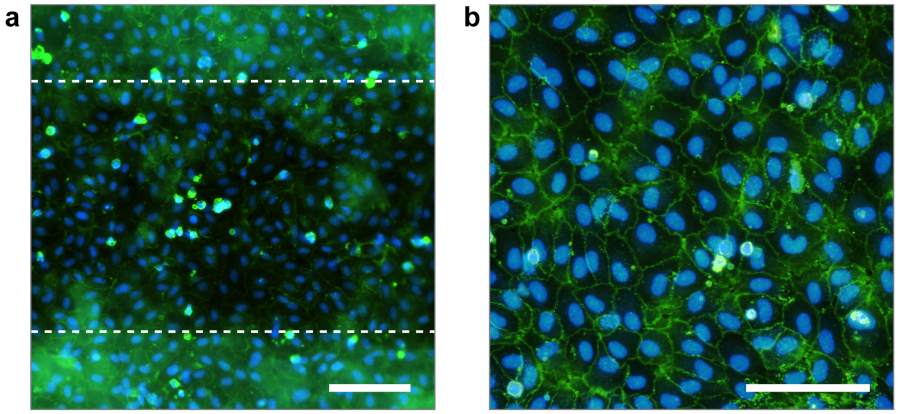


**Supporting Figure 6. Human umbilical vein endothelial cells (HUVECs) formed a continuous cobblestone-like monolayer spanning the entire interface.** (a) Cells were cultured for 7 days on PEDOT:PSS/PEGDE printed on collagen. PEDOT:PSS/PEGDE on collagen region centered and indicated by dotted lines, areas above and below are collagen only regions. (b) Representative image of HUVECs on PEDOT:PSS/PEGDE, showing characteristic cobblestone morphology. Samples were fixed and stained for membrane (green, Wheat Germ Agglutinin) and nuclei (blue, Hoechst). Representative images. Scale bars = 100 µm.

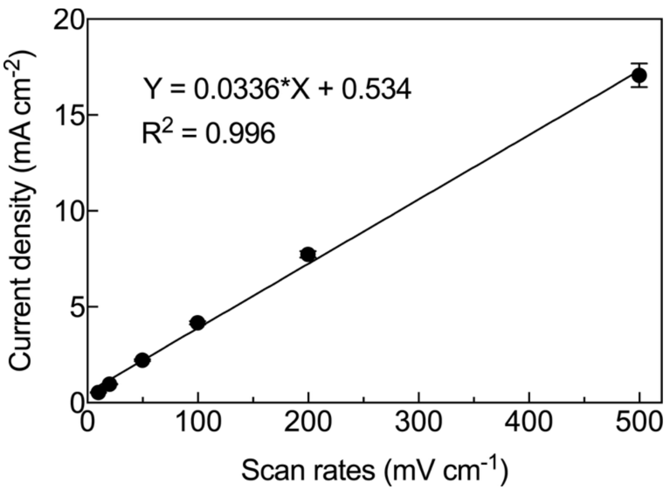


**Supporting Figure 7. Current density of PEDOT:PSS/PEGDE electrodes on collagen versus scan rate.** The plot showed a linear relationship ($R^2 = 0.996$, line represents linear regression fitting), consistent with ideal double-layer capacitive behavior without diffusion limitations. Mean and standard deviation presented. N = 5.

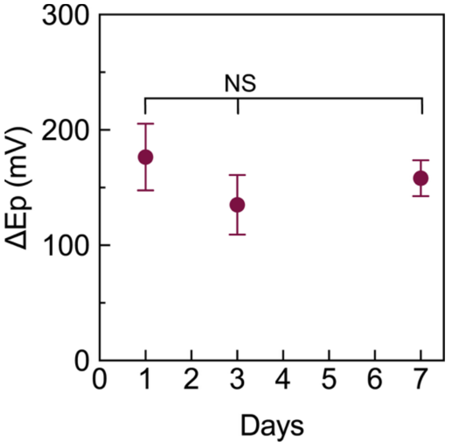


**Supporting Figure 8. Separation between anodic and cathodic peaks (ΔEp) in cyclic voltammetry was not significantly different over 7 days of incubation in DMEM, indicating sustained redox reversibility of collagen-based electrodes under cell culture conditions.** Mean and standard deviation presented. N = 3. One-way analysis of variance (ANOVA, F = 2.21, P = 0.191) and Tukey's multiple comparison tests performed. Non-significant (NS) P > 0.05.

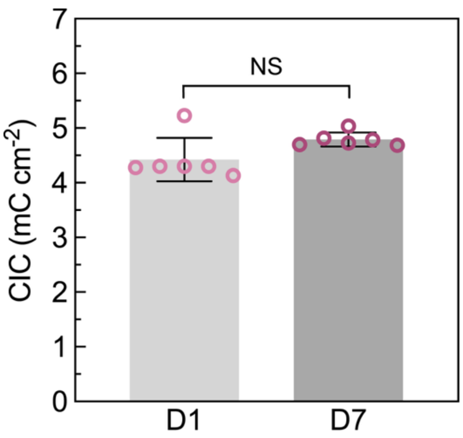


**Supporting Figure 9. Charge injection capacity (CIC) generated by biphasic pulses (±0.5 V vs Ag/AgCl, 1 s) remained unchanged after 7 days in for PEDOT:PSS/PEGDE electrodes on collagen incubated in DMEM.** Separate samples were measured at each time point. N=6. Mean and standard deviation presented. Plot points denote individual samples. Unpaired t-test ($F = 9.45$, $P = 0.0577$). Non-significant (NS) $P > 0.05$.
